# Beyond species-level planning: The role of bioclimatic variation within species distributions

**DOI:** 10.1101/2025.10.27.684983

**Authors:** Thiago Cavalcante, Marta Cimatti, Sara Si-Moussi, Wilfried Thuiller, Moreno Di Marco, Heini Kujala

## Abstract

Conserving biodiversity under a changing climate is a complex challenge that requires comprehensive conservation planning approaches accounting for both current biodiversity patterns and the diverse ecological and environmental changes that species and ecosystems are likely to encounter over time. Systematic conservation planning (SCP) offers a strategic framework to meet this challenge by prioritizing areas that promote species persistence and ecological resilience. Traditionally, SCP focuses on conserving adequate amounts of species’ distributions to ensure their long-term persistence. More recently, partitioning species’ distributions into bioclimatic components has emerged to explicitly represent niche variability, enhancing adaptive capacity by preserving local adaptations and genetic diversity across environmental gradients. Despite this conceptual progress, empirical comparisons of species-level and bioclimatic component prioritization remain scarce. This study aimed to compare species-level and bioclimatic component prioritization by assessing their trade-offs and effectiveness in supporting species persistence and ecological resilience. Specifically, we aimed to (i) assess the surrogacy between species-level and bioclimatic component prioritizations, (ii) examine their spatial overlap and divergence, and (iii) quantify and compare environmental heterogeneity within priority areas identified by each approach. We found that species-level and bioclimatic component prioritizations act as reasonable surrogates for one another overall, but species-level prioritization tended to underrepresent the least-covered bioclimatic components, with failures to capture certain components in the top-ranked areas. Spatial overlap between the two approaches was generally high, though it declined with more restrictive thresholds and under future conditions. Additionally, bioclimatic component prioritizations consistently captured higher within-group multivariate dispersion in environmental heterogeneity in selected areas. Our findings highlight that bioclimatic component prioritization captures greater environmental heterogeneity and complements species-based approaches by better representing niche diversity. Integrating both strategies may offer a more robust path toward climate-resilient conservation planning that accounts for ecological requirements and environmental variation.

## Introduction

Conserving biodiversity under a changing climate is a complex challenge that requires accounting for the many ecological and environmental factors influencing species and ecosystems over time. It requires comprehensive conservation planning approaches that take into account not only current biodiversity patterns but also the diverse range of environmental changes that species and ecosystems are likely to encounter in the future (Garcia et al., 2014b; Pecl et al., 2017; IPCC, 2023). These shifts are rapidly altering natural environments, often in unpredictable ways, with profound implications for biodiversity (IPCC, 2023). To be effective, conservation planning must therefore be dynamic and forward-looking, integrating ecological understanding with predictive models to anticipate and adapt to future changes (Garcia et al., 2014a; Garcia et al., 2014b; Carroll et al., 2015; Pollock et al., 2020). Systematic conservation planning (SCP) offers a strategic framework to navigate these challenges, enabling conservation efforts that are both flexible and effective in safeguarding biodiversity amid changing environmental conditions (Margules and Pressey, 2000; Araújo, 2012; Chauvier-Mendes et al., 2024).

The most common strategy in SCP has traditionally focused on conserving adequate amounts of species’ distributions to ensure their long-term persistence (Margules and Pressey, 2000). This approach aims to protect as much of a species’ geographic range as possible, under the assumption that conserving these areas will safeguard the habitats and ecological requirements necessary for population viability (Wolff et al., 2023). It treats all parts of a species’ range as equivalent, without considering differences in environmental conditions across the range. More recently, an alternative strategy has been proposed: subdividing species’ distributions into bioclimatic components, which are portions of a species’ range defined by distinct climatic conditions (Hanson et al., 2020; Cimatti et al., 2025). The overarching rationale of this approach is to explicitly represent the full variability of species’ niches, that may influence their adaptive capacity and resilience to environmental change (Hanson et al., 2017; Hanson et al., 2020). By capturing distinct climatic environments within a species’ range, this strategy seeks to preserve populations exposed to different selective pressures, which are conditions that can foster local adaptations and maintain intraspecific genetic diversity (Hanson et al., 2017).

Despite this conceptual advancement of incorporating bioclimatic components, there remains limited empirical understanding of how well these approaches perform in practice within spatial conservation prioritization. For many vertebrate species globally, protected areas fail to capture the full range of climatic conditions within their distributions (niche representation), highlighting a critical shortfall in current conservation networks (Hanson et al., 2020). A recent study has also shown that conserving the full range of species’ climatic niche variability can involve trade-offs with the protection of climatically stable areas (often referred to as refugia), which are traditionally prioritized for long-term persistence (Cimatti et al., 2025). While these studies highlight the urgency of expanding protected areas to encompass diverse climatic conditions and demonstrate what this implies in terms of risk prevention, the overall performance of alternative prioritization schemes has not yet been evaluated. In particular, there remains a gap in understanding the extent to which prioritizing bioclimatic components affects species coverage and vice versa, as well as how these approaches differ in effectiveness, spatial agreement, and representation of environmental heterogeneity. These differences are especially relevant in the context of SCP, where accounting for environmental heterogeneity (i.e., the variation in habitat conditions across landscapes) is a critical factor for designing effective protected area networks (Opdam and Wascher, 2004; Araújo, 2012; Carroll et al., 2017; Wang et al., 2018; Wu et al., 2025).

Environmental heterogeneity has long served as a surrogate for biodiversity, under the assumption that areas with greater heterogeneity are more likely to support a higher number of species or diverse ecological functions (Faith and Walker, 1996; Beier and de Albuquerque, 2015). This approach has guided the selection of priority areas by identifying regions that encompass a range of environmental conditions, thereby increasing the likelihood of protecting species’ ecological requirements. By spanning diverse environments, heterogeneity represents a proxy for adaptive genetic diversity, promoting variation that underpins evolutionary resilience and enabling species across broad environmental spectra to adjust locally to changing conditions rather than shift their ranges (Sgrò et al., 2011; Carvalho et al., 2017; Hanson et al., 2017; Hällfors et al., 2024). However, while heterogeneity has long been recognized as a key factor in effective conservation design, it is less clear how different prioritization strategies, such as species-level versus bioclimatic component approaches, capture this variability across landscapes.

Understanding these differences is therefore essential for evaluating how prioritization strategies promote species persistence and ecological resilience under changing conditions. This is particularly important because bioclimatic component approaches primarily reflect climatic variation, whereas environmental heterogeneity encompasses broader habitat conditions (e.g., topography, soil, vegetation structure) that can influence species distributions and adaptive potential.

The main objective of this study was to assess the trade-offs and compare the effectiveness of species-level and bioclimatic component spatial prioritization in identifying areas that support species persistence and ecological resilience. Specifically, we aimed to (i) assess how well species-level prioritization represents species’ bioclimatic components, and vice-versa, through a surrogacy analysis; (ii) evaluate the spatial overlap and divergence between the two prioritization strategies; and (iii) quantify and compare the environmental heterogeneity within the priority areas identified by species-level and bioclimatic component prioritization approaches.

## Methods

### Study system and Datasets

Our study focused on 1,050 terrestrial vertebrate species across four taxonomic classes in Europe: Amphibians (84 species), Birds (501), Mammals (275), and Reptiles (190). These species represent a substantial proportion of those listed under the EU Habitats Directive (Article 17), the Birds Directive (Article 12), and the IUCN Red List of Threatened Species. The study area encompassed the boundaries of the Europe-wide biogeographical regions as defined by the Habitats Directive and the EMERALD Network (EEA, 2024). The two main datasets underpinning the analysis were species distribution models (SDMs), generated using an ensemble of machine learning algorithms (Si-moussi and Thuiller, 2024), and bioclimatic components derived from these SDMs (Cimatti et al., 2025). These datasets were available at a spatial resolution of 1 km^2^.

Species Distribution Models used in this study were generated in a previous analysis using an ensemble of three machine learning algorithms: Random Forest, XGBoost, and multi-layer perceptron neural networks (Si-moussi and Thuiller, 2024). Hyperparameters were optimized using GridSearch in scikit-learn, and class-weighting was applied to correct for presence–absence imbalance, particularly for rare species (see Supplementary Material for additional information on the methodology). Species predictions were constrained using expert range maps (IUCN, BirdLife), with exponential decay smoothing from range edges to buffer zones by taxonomic class. Future distributions were projected for 2041–2070 under the SSP3-RCP7.0 climate and land-use scenario, using three global climate models (GFDL-ESM4, MRI-ESM2-0, UKESM1-0-LL).

Bioclimatic components capturing climatic variability within species’ distributions were obtained from Cimatti et al. (2025). This study applied a two-step clustering framework using the same six ecologically relevant bioclimatic variables that underpinned the SDMs. First, Self-Organizing Maps (SOMs) reduced multidimensional climate data into representative nodes. Then, k-means clustering grouped these nodes into species-specific bioclimatic clusters. To map these clusters back onto both present and future species distributions, a K-Nearest Neighbours (KNN) classifier was trained and validated through 10-fold cross-validation, assigning each distribution pixel to its most probable cluster (see Cimatti et al., 2025 for further methodological details). The outcome was a set of bioclimatic component maps for each species, capturing species-specific climatic variation within both present and future distributions. The bioclimatic component maps for each species were informed by SDM-derived suitability values, so that each component represents the predicted habitat suitability of its locations.

### Spatial prioritisation analysis

We conducted the spatial conservation prioritisation analysis using the Zonation 5 v2.1 software (Moilanen et al., 2022). Zonation generates a hierarchical ranking of importance for each grid cell within the landscape, from most important (1) to least important (0), based on occurrence levels (represented here by habitat suitability) in both present and future distribution layers. The prioritisation process in Zonation is driven by a meta-algorithm that operates based on a marginal loss rule (see Moilanen et al., 2022 for the mathematical formulation). The meta-algorithm determines the ranking (ordering) of grid cells, while the marginal loss rule, operating within the meta-algorithm, calculates how much each feature (i.e., species or bioclimatic components) is reduced when a cell is removed, aggregating these reductions into a cell-specific value. This allows grid cells to be compared, ranked, and ordered in a way that effectively maximizes the representation of each feature in the highest-ranked cells. We used the default CAZ2 rule for marginal loss calculation. CAZ2 slightly reduces the average coverage of features in the top-ranked cells to improve the representation of the least well-represented features, thereby providing a balanced solution across all features included in the analysis. We performed two spatial prioritization strategies: one focusing on species and another on bioclimatic components (feature groups). For each strategy, we conducted separate prioritizations using present and future distribution data (time steps). This resulted in four prioritization variants (2 feature groups x 2 time steps).

### Surrogacy analysis

To compare species-level and bioclimatic component prioritization, we first performed a surrogacy analysis. This means that each time we ran Zonation, we included both species and bioclimatic components in the data but allowed only one group to influence the prioritization outcome. More specifically, we set the weight of the non-surrogate feature to zero so it would not influence the prioritization but would still be included for monitoring. Meanwhile, the surrogate features kept their default weight of 1.0, ensuring that they fully guided the prioritization process. The same procedure was repeated for the present and future. We then generated separate performance curves, which show the cumulative fraction of each feature’s full distribution that would be covered, if grid cells were protected in their optimised priority ranking order. From the individual performance curves, we produced average curves for both surrogate and non-surrogate features, allowing us to assess how well the non-surrogate features are represented in the prioritization based on the surrogate features. Species and bioclimatic components were used sequentially as surrogates and non-surrogates.

### Spatial Overlap

We compared the priority maps produced by species-level and bioclimatic component prioritizations to evaluate how much they overlap and differ spatially. We used two complementary metrics for this comparison: Schoener’s D and the Jaccard index. Schoener’s D is a metric for continuous values that captures the degree of spatial similarity between the maps. It is calculated by normalizing the values in each map, calculating the absolute differences between corresponding values, and summing these differences. The result is scaled to give a value between 0 (no overlap) and 1 (complete overlap), reflecting broader distributional similarities.

To specifically compare high-priority areas, we binarized the maps by selecting the top fraction of ranked cells (the top 30%). For these binary comparisons, we used the Jaccard index, which quantifies overlap by measuring the ratio of shared priority areas relative to the total areas in both maps, thus reflecting the extent of agreement between species-level and bioclimatic prioritization at a specific threshold. Together, Schoener’s D and the Jaccard index provide complementary perspectives, with Schoener’s D capturing overall similarity across the continuous priority maps and the Jaccard index focusing on overlap in the top-ranked priority areas.

### Environmental heterogeneity

We measured environmental heterogeneity using spectral heterogeneity derived from satellite imagery, which captures variation in surface reflectance values across the landscape (Rocchini et al., 2010b; Torresani et al., 2024). We used the MODIS surface reflectance data product (MOD09A1, Collection 6.1; available at https://www.earthdata.nasa.gov/data/catalog/lpcloud-mod09a1-061) to characterize spatiotemporal patterns of surface reflectance across the study area. This product provides atmospherically corrected 8-day composites of seven land surface reflectance bands at 500 m spatial resolution (Vermote, 2015). The MOD09A1 image collection was filtered to include scenes acquired between January 1, 2014, and December 31, 2024. To ensure high-quality observations, we applied a Quality Assurance (QA) filtering procedure based on the StateQA bitmask. Poor-quality observations contaminated by clouds, cloud shadows, adjacency to clouds, snow, and fire were excluded. Spectrally, we limited our analysis to bands most commonly used in land surface studies: band 3 (Blue), band 4 (Green), band 1 (Red), and band 2 (Near Infrared). After QA filtering and band selection, we calculated the temporal standard deviation of surface reflectance for each band across the full study period.

We then conducted a multivariate analysis to quantify and compare spectral heterogeneity between species-level and bioclimatic component-level prioritization. We extracted reflectance band values within the top 10% priority areas identified by each approach, as this threshold captures greater spatial divergence between them. The extracted values were scaled and used to calculate multivariate dispersion (betadisper) based on Euclidean distances. Because calculating pairwise distance matrices requires comparing all observation pairs, causing computation time to grow quadratically with sample size [O(n^2^)], we implemented a subsampling strategy using random subsets of 1,000 planning units from the full dataset. We performed 100 independent replicates of the analysis on these subsets to quantify the variability and uncertainty introduced by subsampling. This method balances computational feasibility with robustness, enabling us to capture consistent patterns in multivariate dispersion while explicitly accounting for sampling variability in the results. We used the *betadisper()* function, implemented in the *vegan* R package (Oksanen et al., 2025), to assess the multivariate homogeneity of group dispersions by calculating the average distance of each observation to the group centroid in multivariate space. This provides a measure of within-group variance (dispersion) in spectral heterogeneity, indicating how much more heterogeneity there is within each prioritization approach. To assess whether differences in dispersion between groups were statistically significant, we used a permutation test via *permutest*.*betadisper()*, which permutes model residuals to generate a distribution of the F-statistic under the null hypothesis of equal group dispersions. We used 999 permutations in our analysis. Spectral heterogeneity was evaluated exclusively for present period prioritizations, ensuring the environmental variability assessment is based on empirically observed conditions. All data processing and statistical analyses were conducted using Google Earth Engine (Gorelick et al., 2017) and R (v. 4.4.3) (R Core Team, 2025).

## Results

We found that both the species and bioclimatic components are good surrogates for each other based on how well they represent the non-surrogate group on average (Figure 1). However, prioritization at the species level resulted in a poorer performance for the bioclimatic components in the sense that some individual bioclimatic components were missed entirely from the top-ranked grid cells (minimum performance curves, Figure 1).

**Figure 1.**
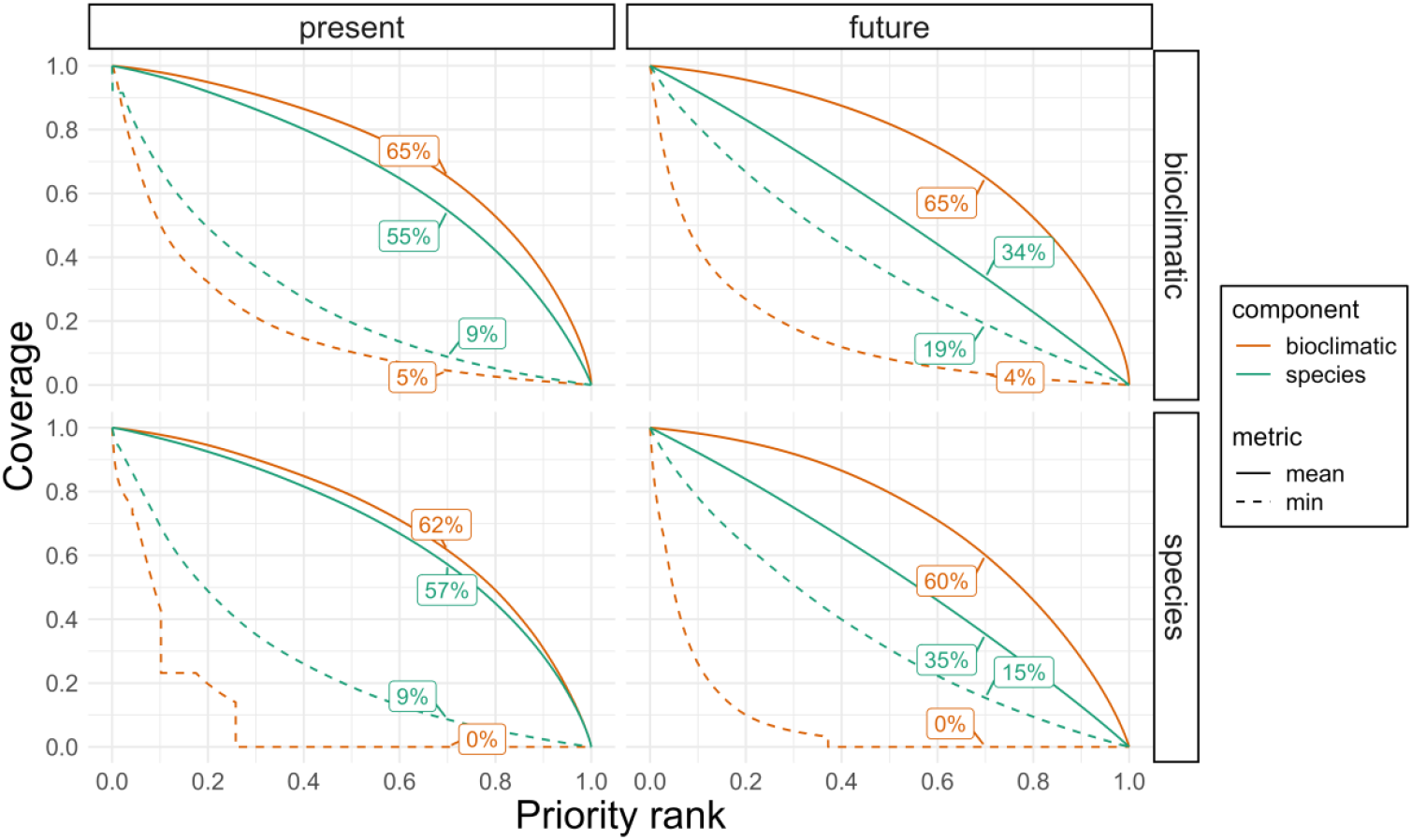
Coverage of species and bioclimatic components across prioritization ranks in present and future scenarios, based on the surrogacy analyses. Coverage values at the top 30% of the priority ranking are indicated by percentage labels. Facets are organized by features (bioclimatic vs. species) and planning period (present vs. future), with features in each row serving as the surrogate, while the opposite component is tracked as a non-surrogate.

Despite some bioclimatic components being missed, a considerable proportion of them remained represented within top-priority areas. For instance, when selecting the top 30% of the landscape in species-level prioritizations, only 14 species in the present and 3 in the future had any associated bioclimatic component with zero representation. However, these numbers increased to 46 in the present and 52 in the future when considering bioclimatic components with less than 5% coverage. When examining the overall coverage distributions within the same top 30%, the bioclimatic and species datasets appeared broadly similar in the present, with noticeable differences only at very high coverage values (Figure 2). In contrast, future prioritizations showed more pronounced divergence with species-level coverage showing high homogeneity across species at low to intermediate values, and bioclimatic data exhibiting high heterogeneity of coverage.

**Figure 2.**
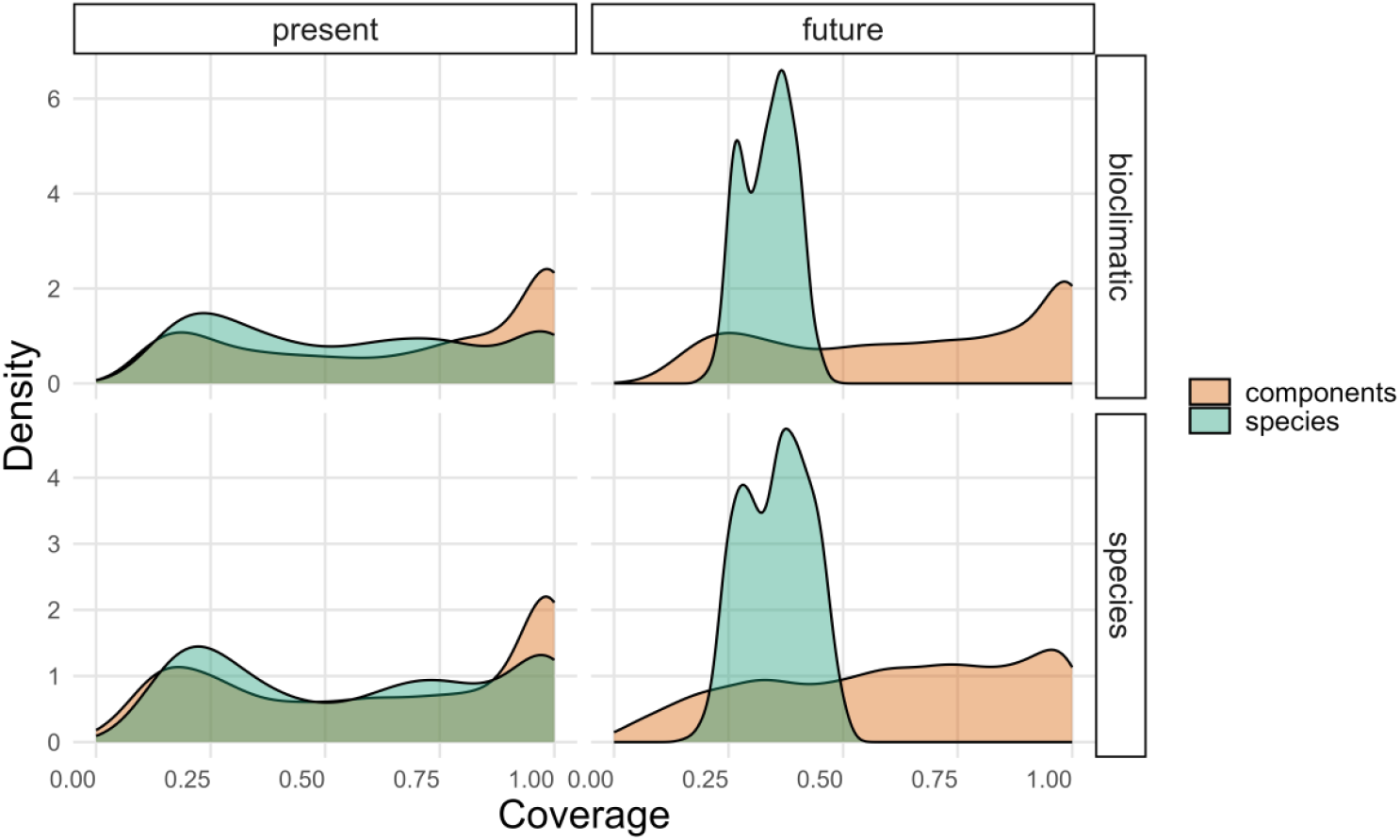
Density plots comparing the coverage distributions of species-level and bioclimatic component-level prioritizations for the present and future in the top 30% priority areas. Kernel density estimates are shown to provide a smoothed representation of the underlying distribution. The x-axis shows the actual coverage range and the y-axis indicates relative density.

The priority areas identified by species-level and bioclimatic component-level strategies exhibited considerable spatial similarity overall (Figure 3).

**Figure 3.**
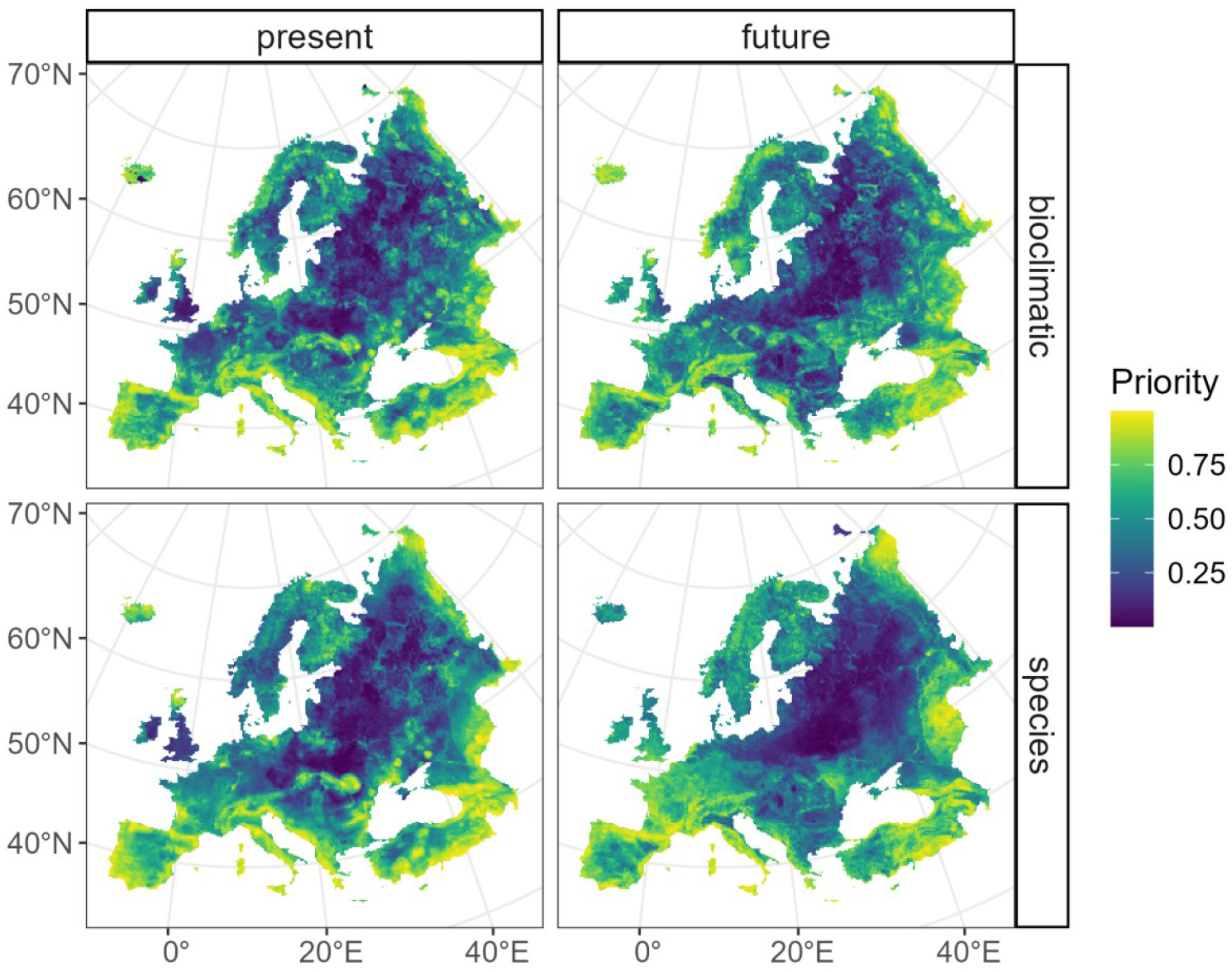
Spatial patterns of priority conservation areas under present and future climate scenarios, showing species-level and bioclimatic component-level prioritizations. Priority values range from 0 (lowest) to 1 (highest).

According to Schoener’s D, a measure of the similarity of continuous prioritization values across the landscape, spatial congruence was high in the present (D = 0.90) but exhibited a slight reduction in the future (D = 0.86). However, when comparing binary representations of the top 30% priority areas, the Jaccard index showed an overlap of 0.645 for the present, which dropped to 0.517 in the future, highlighting areas of significant divergence (Figure 4). For the more restrictive top 10% priority areas, overlap was even lower, with a Jaccard index of 0.448 for the present and 0.313 in the future, revealing greater divergence at higher prioritization thresholds (Table 1).

**Table 1.**
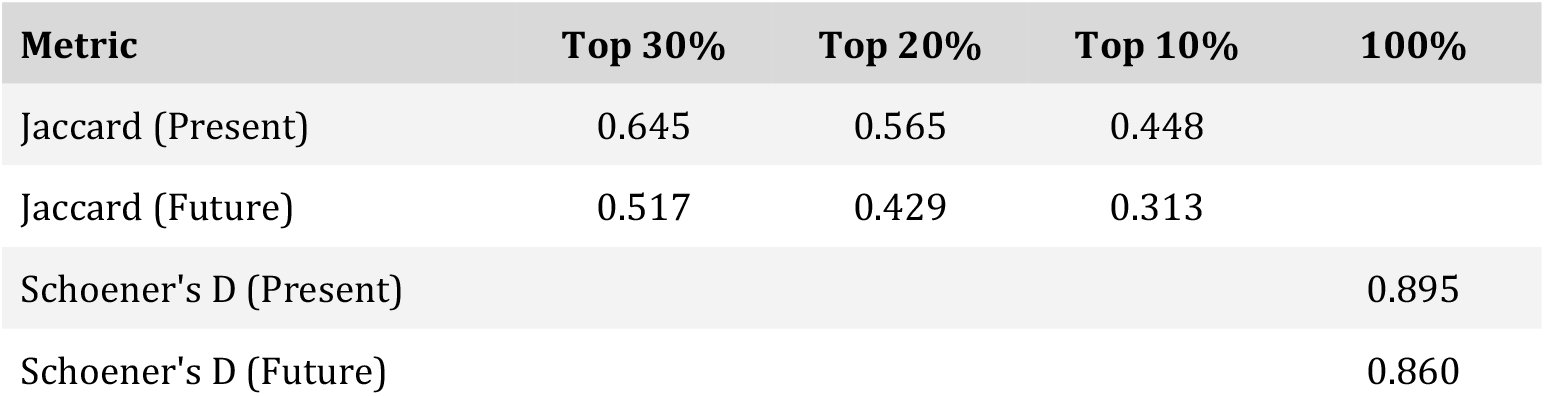
Overlap metrics between species-level and bioclimatic rank maps under present and future conditions. Jaccard indices are calculated for the top 30%, 20%, and 10% priority areas. Schoener’s D represents continuous overlap across the entire range (100%).

**Figure 4.**
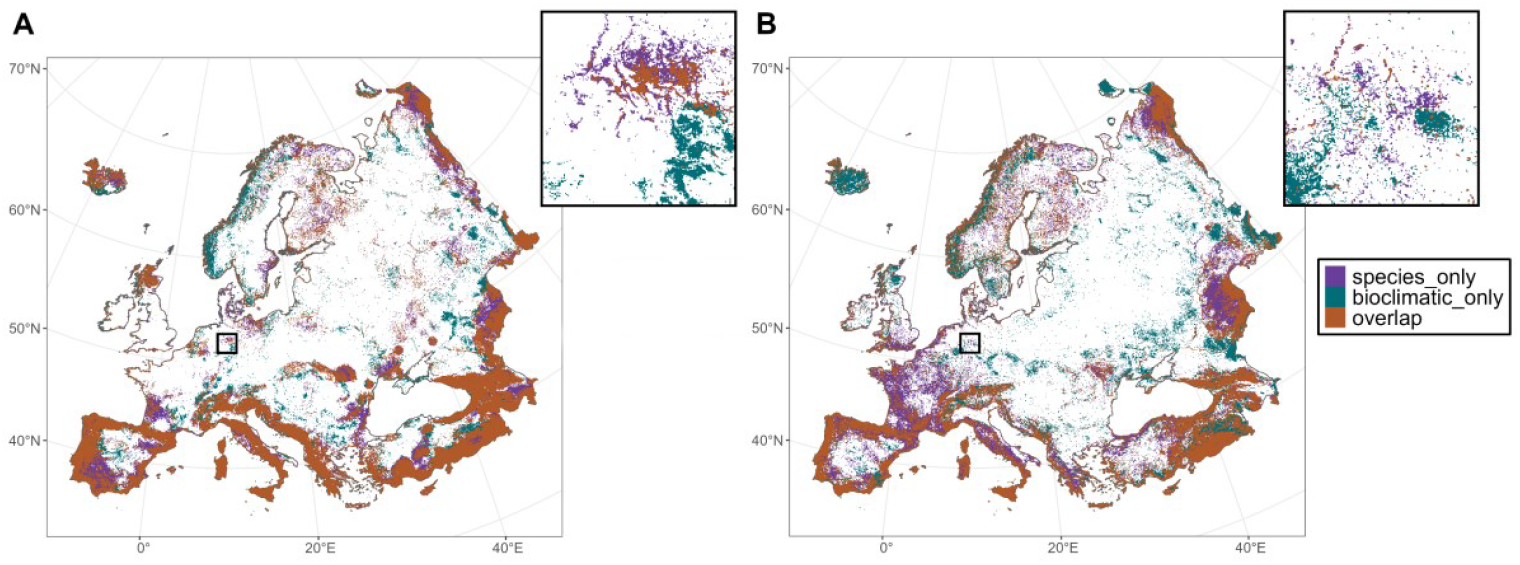
Spatial patterns of overlap and divergence between species-level and bioclimatic component-level strategies within the top 30% priority areas under present (A) and future (B) climate scenarios. Insets provide zoomed views of selected regions to highlight spatial details.

We found that bioclimatic component-level prioritization areas showed higher temporal variability, as indicated by band-wise differences in the standard deviation of reflectance values (Figure 5A). This means that these areas experienced more changes in environmental conditions over time, as detected by the satellite data. Spectral heterogeneity, measured as multivariate dispersion, was consistently higher in bioclimatic component-level prioritization areas (mean dispersion = 1.70, SD = 0.048) compared to species-level prioritization areas (mean dispersion = 1.13, SD = 0.075) across 100 replicates (Figure 5B and 5C). The range of dispersion values was also narrower for bioclimatic prioritization (1.61–1.82) than for species prioritization (0.95–1.32). Permutation tests showed highly significant differences in multivariate dispersion between groups across all replicates (p ≤ 0.001), indicating consistent differences in spectral heterogeneity.

**Figure 5.**
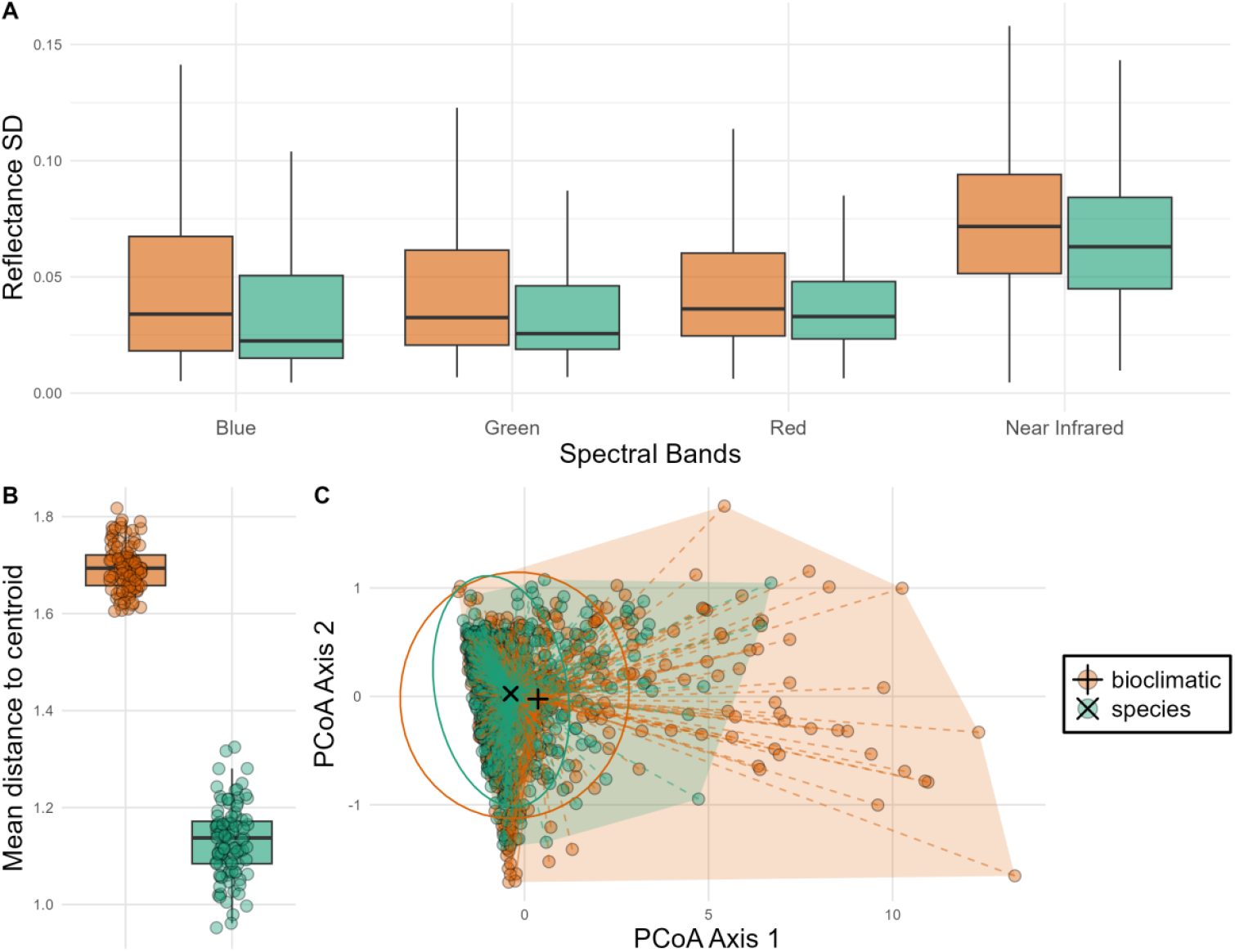
(A) Temporal variation in reflectance values (standard deviation, SD) across spectral bands. (B) Within-group multivariate dispersion (average distance to centroid) across 100 replicates, comparing bioclimatic component-level and species-level prioritization. Individual replicate values are shown as jittered points. (C) Principal coordinates analysis (PCoA), based on one replicate, illustrating the multivariate dispersion of spectral data by component. Points represent planning unit sites, with dashed segments connecting each site to its group centroid (black shapes). Convex hull polygons and 95% confidence ellipses illustrate the spatial extent and variation within groups.

## Discussion

This study aimed to compare species-level and bioclimatic components prioritization approaches by examining their effectiveness, spatial agreement, and representation of environmental heterogeneity. While both approaches reflected similar patterns in priority areas overall, we found that notable asymmetries and trade-offs emerge, particularly when considering the spatial divergence at higher prioritization thresholds and under the future climate change scenario.

Species-based prioritization was less effective at representing the full range of climatic conditions compared to a bioclimatic-based approach, demonstrating the risk of disregarding niche variability when performing a prioritization at species range level (Hanson et al., 2020; Cimatti et al., 2025). While species-level priorities covered a broad range of bioclimatic components overall, the worst-off features (i.e., those with the lowest remaining representation because they occur in small areas or have low redundancy) were left out early in the ranking process. In contrast, bioclimatic-based prioritization maintained a more balanced representation, ensuring that components with limited extent or low redundancy were preserved throughout the prioritization. Interestingly, despite not using species data directly, the bioclimatic approach still achieved similar levels of mean and minimum species representation compared to the species-based prioritization. These findings are consistent with recent studies highlighting widespread shortcomings in the representation of environmental niches within protected areas (Hanson et al., 2020), which can impact species’ ability to adapt to environmental change (Moritz, 2002; Sgrò et al., 2011; Hanson et al., 2017). Beyond emphasizing the importance of capturing environmental conditions, our results go further to demonstrate that preserving these climatic niches can simultaneously serve as a fair and representative surrogate for species-level conservation. This proves effective not only for broadly distributed species but also for the worst-off features, as reflected by higher values along the minimum curves throughout the prioritization (Figure 1).

When examining coverage distributions of species-level and bioclimatic components approaches within the top 30% priority areas, we found broadly similar patterns under present conditions, in agreement with the average performance curves. However, the divergence observed in the future scenario was striking, with species-level coverage peaking at relatively low to intermediate levels. These patterns revealed by the coverage results were mirrored in the spatial configuration of priority areas. Overall, we found strong spatial convergence in prioritization across the landscape, yet notable divergence emerged within the top priority areas, with divergence intensifying under the future climate scenario. This may be partly attributed to common issues of overprediction in SDMs under climate change scenarios (Thuiller, 2004; Guisan and Thuiller, 2005). Future projections often identify broad areas of habitat suitability, potentially inflating the number of grid cells where species are predicted to occur. This can lead to prioritizations that assign high priority ranks to regions where species do not occur and/or underestimate the effectiveness of selected priority areas (i.e., the proportion of species’ ranges covered) (Velazco et al., 2020). In contrast, the climatic profiles identified through bioclimatic clustering and projected into future scenarios may be less prone to such modeling artifacts. This is primarily because bioclimatic components summarize broader, regional environmental conditions rather than the often-complex ecological niches of individual species. In other words, bioclimatic clustering summarizes the core climatic components within species’ ranges without trying to fit the complex response curves used in SDMs. As a result, future projections based on climatic clustering can provide a conservative framework for spatial prioritization under climate change when species-level dispersal constraints and biotic interactions are uncertain or unmodeled.

We found that spectral heterogeneity, measured by within-group multivariate dispersion (i.e., the average distance to the group centroid), was consistently higher in areas selected through bioclimatic prioritization across all replicates. These areas also exhibited higher temporal standard deviation, reflecting greater environmental variability across space and time (Figures 5). Such heterogeneity may function as a climatic buffer, offering a diversity of conditions that could enhance the adaptive capacity of species facing climate change (Jetz and Rahbek, 2002; Ohlemüller et al., 2008; Wu et al., 2025). This is notable from the perspective that variation in surface reflectance values across the landscape captures fine-scale habitat complexity and structural variation (e.g., land cover diversity, vegetation types, canopy structure) that are often overlooked by bioclimatic variables (Rocchini et al., 2010b; Tuanmu and Jetz, 2015; Randin et al., 2020; Torresani et al., 2024). Such heterogeneity is associated with a broader diversity of ecological niches (Gould, 2000; Rocchini et al., 2010a; Torresani et al., 2024), which in turn supports species adaptation and long-term persistence. However, while bioclimatic approaches perform well in representing environmental heterogeneity, they do not incorporate the direct species–environment relationships inherent to SDMs. By prioritizing climatic gradients rather than modeled species niches, bioclimatic clustering may include areas of low habitat suitability or those outside core species ranges, such as sink populations or marginal habitats, that prioritization based on SDMs tends to avoid. This highlights an important trade-off: maximizing environmental heterogeneity can support broad-scale resilience, but it may do so at the expense of focusing conservation efforts on high-quality habitats essential for individual species’ persistence. In our study, we addressed this trade-off by incorporating SDM-derived suitability values into the bioclimatic components, thereby grounding the climatic clusters in biologically meaningful habitat information.

The differences between approaches also stem from their distinct assumptions and strengths. Bioclimatic clustering is based on the premise that climatic variability underpins adaptive potential (Hanson et al., 2017; Hanson et al., 2020). It assumes that protecting a wide range of climatic conditions within a species’ range helps maintain local adaptations and that species have the capacity to respond to environmental changes through adaptation. In this view, bioclimatic components serve as proxies for intraspecific variation, especially in contexts where genetic and population data are not spatially comprehensive enough (Hanson et al., 2017). In contrast, SDMs rely on empirically observed relationships between species occurrences and environmental predictors, producing spatially explicit projections grounded in the strong theoretical framework of niche theory (Soberon and Peterson, 2005; Soberón and Nakamura, 2009). However, they also rest on the important assumption that a species’ response to environmental conditions will remain consistent over time (Soberón and Nakamura, 2009). Taken together, these differences highlight that the two approaches are not mutually exclusive but offer complementary insights: while SDMs highlight core habitats critical for long-term persistence, bioclimatic components reflect the environmental heterogeneity that may underpin species’ adaptive responses.

Our results should be interpreted with an awareness of certain methodological nuances. In this study, the bioclimatic component prioritization was informed by continuous habitat suitability values derived from SDMs to ensure that the bioclimatic components reflected biologically meaningful habitat quality. This means that while the prioritization approaches differ, with one focusing on the species’ entire distribution and the other on their climatic clusters, they are not based on independent datasets. Rather, both approaches rely on the same underlying SDM data, partitioned in different ways. We would expect that using thresholded binary maps or models derived from independent data could result in more pronounced differences, potentially revealing sharper trade-offs between the two prioritization strategies. More broadly, the degree to which species-based and bioclimatic-based prioritizations act as effective surrogates for each other and/or contribute to capturing environmental heterogeneity will be inherently dependent on the input data. Differences in taxonomic resolution, spatial coverage, and climatic variability can substantially influence how well one approach represents the other. For instance, in regions with high environmental heterogeneity but limited species data available for building SDMs, such as mountainous or less-sampled tropical areas, bioclimatic components may capture environmental heterogeneity more effectively than species-based models derived from sparse or biased data. However, when reliable data are available, SDMs offer the added benefit of explicitly modeling the ecological relationships that shape species distributions, enabling prioritizations that target the most suitable and ecologically meaningful areas within a species’ range—something that bioclimatic components alone cannot capture.

Looking ahead, combining species’ ecological requirements with environmental surrogates through integrated prioritization strategies could offer promising avenues to improve spatial conservation planning under climate change (Buenafe et al., 2025). Such approaches recognize the need to balance multiple, sometimes conflicting, conservation objectives to enhance resilience in response to emerging environmental challenges (e.g., Cimatti et al., 2025). In our study, we took a step in this direction by incorporating SDM-derived suitability values into the bioclimatic component prioritization process. More broadly, our results demonstrate several key insights: (i) bioclimatic component prioritization offers a more balanced representation of climatic gradients and performs well as a surrogate for species representation; (ii) areas selected through bioclimatic prioritization consistently exhibited higher environmental heterogeneity, which is crucial for supporting species adaptation and resilience; (iii) species-based prioritization remains essential for targeting high-quality habitats but may miss rare or underrepresented climatic conditions critical for long-term resilience; and (iv) important trade-offs and divergences emerge between approaches, particularly under future climate scenarios. Together, these findings underscore that no single method is universally superior, and that hybrid strategies integrating ecological specificity and environmental heterogeneity can lead to more adaptive, robust, and climate-resilient outcomes in systematic conservation planning.

## Acknowledgments

This research was supported by the NaturaConnect project, funded by the European Union’s Horizon Europe research and innovation programme (grant agreement No. 101060429).

The authors acknowledge CSC – IT Center for Science, Finland, for providing access to high-performance computing resources.

