## Supplementary Material for "Beyond species-level planning: The role of bioclimatic variation within species distributions"

**Affiliations:**
[1] Finnish Museum of Natural History, University of Helsinki, Helsinki, Finland.
[2] Dept. of Biology and Biotechnologies "Charles Darwin", Sapienza University of Rome, Rome, Italy
[3] Univ. Grenoble Alpes, Univ. Savoie Mont Blanc, CNRS, LECA, Grenoble, France

### Species Distribution Models

#### Study extent and target taxa

Species distributions were modelled at 1 km² resolution for terrestrial vertebrates across Europe. The calibration domain extended beyond the **EEA39 reporting region (EU27, the United Kingdom, the four EFTA countries—Iceland, Liechtenstein, Norway, Switzerland—and Turkey)** to include adjacent areas in Russia, North Africa, and the Middle East thereby reducing niche truncation (Thuiller et al., 2004; Barbet-Massin et al., 2010).

The initial vertebrate pool comprised 1,259 species across Amphibia, Aves, Mammalia, and Reptilia from the TetraEU database (Maiorano et al. 2020). This list covers most species reported under the EU Habitats (Article 17) and Birds (Article 12) Directives, as well as many species of conservation concern on the IUCN Red List. After data curation, 1,210 species were retained for modelling.

#### Environmental covariates

Environmental covariates were assembled at 1 km² resolution from publicly available datasets. Climatic variables (temperature, precipitation, snow cover trends) were obtained from CHELSA Karger et al. (2017, 2020). Topographic variables (slope, landforms, coastal proximity) were derived from EarthEnv (Amatulli et al 2018). Soil physical and chemical properties (pH, soil organic carbon, texture) were taken from SoilGrids Poggio et al. (2021). Hydrographic layers describing the extent and salinity of water bodies and wetlands were compiled from various land cover products; CORINE Land Cover 2018, ESA CCI Land Cover and the Global Lakes and Wetlands Database (GLWD) Lehner et al. (2004, 2025), while river density was computed using the EU-Hydro (EEA 2019) and HydroSHEDS Lehner et al. (2008) watercourses polygons. Land systems and land-use intensity were obtained from Sandström et al. (2023). To minimise collinearity, only variables with pairwise correlation < 0.70 were retained.

**Occurrence data and pseudo-absences**

Species observations were obtained from GBIF (1980–2023; location uncertainty < 1 km) between 4–9 January 2023[1]. Records were cleaned with **CoordinateCleaner** (Zizka et al., 2019). Non-avian vertebrates were complemented with IUCN ranges, and birds with breeding ranges from BirdLife International. To account for potential range changes since the last expert assessment, ranges were rasterised and buffered to include 90% of outlier GBIF records within each class. Species with a high proportion of records outside buffered ranges were manually reviewed. Sampling effort maps were generated from observation density per pixel across predefined target groups: Amphibians, Birds, Small Mammals, Large Mammals, Chiroptera and Reptilia.

Pseudo-absences were generated following Barbet-Massin et al. (2012). Three categories were defined: (1) certain presences within buffered ranges, (2) certain absences sampled outside buffered ranges, and (3) uncertain absences sampled within buffered ranges, weighted by target group sampling effort. For 328 species lacking GBIF records but with available expert maps, datasets were generated by sampling certain presence pixels within IUCN ranges proportional to range size and certain absence pixels outside the range. Conversely, for 52 species with GBIF records but no expert range datasets included all observed presences and pseudo-absences. Species with not enough GBIF records (>20 records) and no IUCN ranges were modeled using habitat directives data (n=2) and dropped otherwise (n=49). The procedure was repeated five times per species. Adjustments were made for data-deficient species.

**Species distribution models**

Species distribution models were fitted using Random Forest (RF) Breiman et al. (2001), extreme gradient boosting (XGBoost) Chen et al. (2015), and multilayer perceptron neural networks (MLP) Lecun et al. (2015). Hyperparameters were tuned with GridSearch in scikit-learn Pedregosa et al., (2011). For RF, tree depth and feature subsampling were tuned; for XGBoost, regularisation was additionally optimised; for MLP, architectures of varying depth and width were tested. Class weights were applied to correct for imbalance between presence and absence particularly for rare species.

Presence–pseudoabsence data were partitioned into five spatial blocks (Roberts et al., 2017). Models were trained on four blocks and tested on the fifth. Only models with cross-validated True Skill Statistic (TSS) > 0.4 were retained. Ensembles combined models across algorithms, folds and pseudo-absence replicates.

**Ensemble forecasting**

After calibration and validation, SDMs were projected to maps of environmental suitability ranging from 0 (unsuitable) to 1 (highly suitable). Ensembles were built from all models that met the performance threshold (TSS ≥ 0.4) and combined into a single output: the mean suitability across retained models.

---

[1] Aves : https://doi.org/10.15468/dl.6ptbvk ; https://doi.org/10.15468/dl.6am8ch ; https://doi.org/10.15468/dl.6q3zpq ; https://doi.org/10.15468/dl.y8nfqw ; https://doi.org/10.15468/dl.gdta5n ; https://doi.org/10.15468/dl.tn2b3w ; https://doi.org/10.15468/dl.2bhy85 ;
Amphibia/Mammalia/Reptilia: https://doi.org/10.15468/dl.rxfzz3

**Post-hoc spatial constraints**

Suitability maps were constrained with expert ranges using the **Estar framework** (Hoareau et al., in prep). Post-hoc spatial constraints were applied by clipping suitability maps to IUCN extent-of-occurrence ranges, smoothed with an exponential decay kernel from range boundaries to buffer limits.

**Threshold optimisation procedure**

For each species, thresholds were derived using the Continuous Boyce Index calibration curve (Hirzel et al., 2006). The calibration curve compares the frequency of presences in predicted suitability bins with the frequency expected under random sampling. Ratios > 1 indicate that presences occur more often than expected, ratios ≈ 1 indicate frequencies close to random expectation, and ratios < 1 indicate fewer presences than expected. From this curve, two thresholds (th_low and th_high) were identified to delimit unsuitable, uncertain and suitable probability ranges. In the constrained maps, only pixels with values above th_high were retained, while the remaining were set to zero.

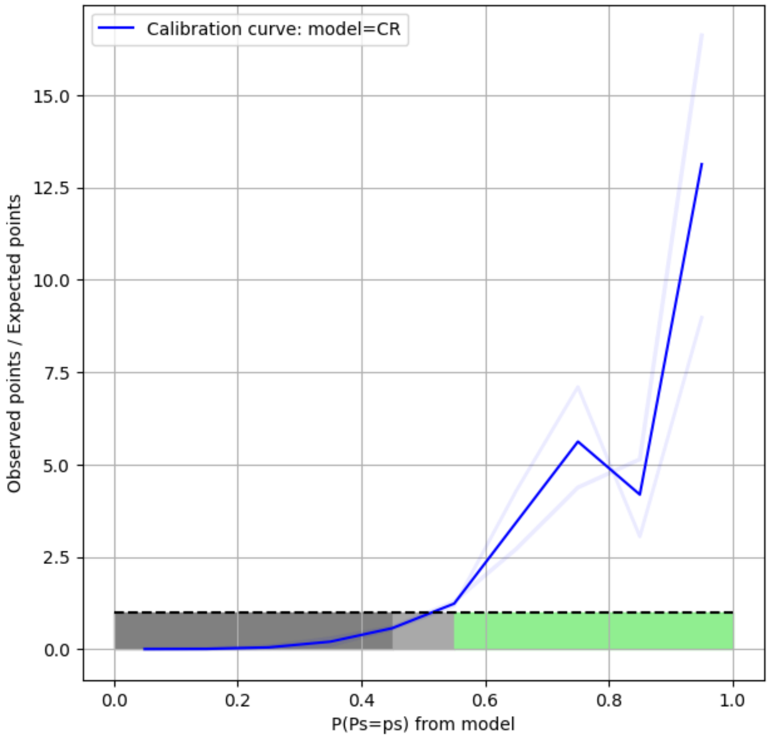


*Figure 1 Example of threshold optimisation using the Boyce Index calibration curve (Hirzel et al., 2005). The curve shows the ratio between the frequency of observed presences in each suitability bin and the frequency expected under random distribution. Ratios < 1 indicate that presences occur less often than expected, ratios ≈ 1 indicate frequencies close to random expectation, and ratios > 1 indicate that presences occur more often than expected. Two thresholds (th_low=0.45 and th_high=0.55) were derived from the curve.*
